# Small Extracellular Vesicle Cargoes Associated with Lymph Node Metastasis in Head and Neck Cancer

**DOI:** 10.64898/2025.12.31.697101

**Authors:** Adnan Shafiq, Shinya Sato, Bahnisikha Barman, Emily N. Arner, Brandee Brown, Alissa M. Weaver

## Abstract

Head and neck squamous cell carcinoma (HNSCC) is a highly aggressive malignancy in which cervical lymph node metastasis critically determines patient prognosis. Despite advances in clinical staging based on tumor size, depth of invasion, and nodal status, these parameters fail to capture the biological heterogeneity of HNSCC, leading to overtreatment or undertreatment, and significant morbidity. Emerging evidence implicates small extracellular vesicles (SEVs) as key mediators of tumor progression and promising biomarkers for metastatic potential. In this study, we performed iTRAQ-based proteomic profiling of SEVs from metastatic (MOC2) and nonmetastatic (MOC1) HNSCC mouse cell lines and identified multiple cargoes associated with metastatic processes, including angiogenesis, extracellular matrix remodeling, immune modulation, and perineural invasion. Further, we employed an immune-competent orthotopic mouse oral carcinoma (MOC) model to investigate how exosome biogenesis affects HNSCC metastasis. Loss of the exosome/SEV biogenesis protein HRS in MOC2 HNSCC cells impaired SEV formation *in vitro* and tumor growth and metastasis *in vivo.* Consistent with the SEV cargoes enriched in MOC2 EVs, immunohistochemical analyses of MOC2 tumors revealed reduced blood vessel and nerve density in HRS-deficient tumors. Analysis of candidate biomarker SEV cargoes in the circulating EVs from HPV-negative HNSCC patients revealed significant correlations of these proteins with metastatic status. Collectively, these findings identify SEV cargoes as potential functional mediators of metastasis and liquid biopsy biomarkers in HNSCC.

## Introduction

Head and neck squamous cell carcinoma (HNSCC) is the sixth most common cancer worldwide, encompassing malignancies arising in the oral cavity, oropharynx, larynx, and hypopharynx (1–3). Among these, oral squamous cell carcinoma (OSCC) accounts for approximately 90% of all HNSCC cancers (2). The prognosis for HNSCC remains poor, particularly for human papillomavirus (HPV)-negative tumors, which are typically associated with greater genomic instability, frequent alterations in tumor suppressor genes, and exposure to environmental carcinogens such as tobacco and alcohol (3–5). While localized HNSCC has a relatively favourable five-year survival rate exceeding 80%, this dramatically decreases to 40% with lymph node (LN) (6) involvement and to approximately 20% when rare distant metastases occur (7).

Cervical LNs are the major metastatic site, being present in 25-35% of patients at diagnosis. Cervical LN metastasis serves as a critical predictor of recurrence, distant disease progression, and overall survival (8, 9). This metastatic process is driven by both tumor-intrinsic and tumor-extrinsic mechanisms, including extracellular matrix (ECM) remodeling, loss of cell-cell adhesion, angiogenesis, neurogenesis, immune evasion, and enhanced migratory and invasive capabilities (3, 4, 6, 10–14).

Tumors can influence the tumor microenvironment (TME) and promote metastasis through the secretion of extracellular mediators (15, 16). Among these, extracellular vesicles (EVs) act as mediators of intercellular communication through their protein, lipid, and nucleic acid cargoes, thereby influencing diverse biological processes including immune modulation, angiogenesis, migration, and invasion (14, 15, 17–19). They are broadly classified into exosomes, which are small EVs (SEVs) (50–150 nm) derived from the endosomal pathway, and ectosomes (50–1000 nm), which form by outward budding of the plasma membrane. SEVs are enriched in tetraspanins (CD9, CD63, CD81), ESCRT-related proteins, and endosomal markers, and they carry a molecular signature reflective of their parent cell, making them particularly relevant for liquid biopsy (17, 20).

In this study, we investigated the role of SEVs in promoting LN metastasis in HNSCC. We employed multiplexed Isobaric Tagging Technology for Relative Quantitation (iTRAQ) mass spectrometry to compare EV protein cargoes from paired nonmetastatic and metastatic MOC HNSCC cell lines. We also employed a well-characterized, immune-competent syngeneic orthotopic mouse model that recapitulates the biological and pathological features of human HPV-negative HNSCC, including its immunosuppressive TME and metastasis to cervical LNs (21). In vivo functional assays revealed that inhibition of exosome secretion with the knockout (KO) of hepatocyte growth factor-regulated tyrosine kinase substrate (HRS) greatly reduced the LN metastasis of MOC2 cells from tongue tumors and moderately reduced tumor size. There was also a decrease in the density of nerves and blood vessels and an increase in CD8^+^ T and CD45^+^ cell penetration into HRS KO MOC2 tumors. We further investigated whether cargoes identified to be enriched in MOC2 SEVs correlate with HNSCC metastasis in patients. Indeed, five candidate proteins, SYT1, STXBP1, TGFβ1, ALCAM, and ITGA5, were enriched in SEVs from HNSCC patients and further correlated with lymph node metastasis status. Together, these data suggest that SEV cargo profiles may reflect metastatic potential and support the development of minimally invasive biomarkers for HNSCC progression.

## Results

### Quantitative proteomic profiling of HNSCC SEVs identifies metastasis-associated protein cargoes

To identify SEV cargoes associated with metastasis, we performed a comprehensive comparative analysis of SEV protein cargos derived from matched mouse oral carcinoma (MOC) cell lines with different metastatic potential. Thus, MOC1 cells are non-metastatic and form “immune-hot” tumors, whereas MOC2 cells are metastatic and form “immune-cold” tumors (21). Differential centrifugation of conditioned media from the two cell lines was performed to remove cells and debris, followed by isolation of large EVs by centrifugation of the supernatant at 10,000 x g for 30 min. SEVs were then purified from that supernatant by density gradient ultracentrifugation (see methods, (22)). Western blot analysis was performed on SEVs purified from the twelve density gradient fractions, along with whole cell lysates (WCL) and large EVs (lEVs) (Supplementary Figure 1A). The blots were probed for classical exosomal markers such as TSG101, CD63 and ALIX, while GM130 served as a negative control to confirm the absence of non-cellular components (Supplementary Figure 1A). While the size and morphological characteristics of MOC1 and MOC2 SEVs were similar, MOC2 cells secrete more SEVs than MOC1 cells (Supplementary Figure 1B-D). We utilized an iTRAQ mass spectrometry approach, which gives a highly quantitative comparative readout of protein abundance (23, 24). A total of 1257 proteins were identified (Supplementary Table 1). Of these, 639 proteins were enriched in MOC2-SEVs and 617 were enriched in MOC1-SEVs (Supplementary Table 1). Because applying a Benjamini–Hochberg (BH) adjusted p-value cutoff excluded many candidates (Supplementary Table 1, statistically significantly enriched candidates highlighted in orange (MOC2) or green (MOC1)), we adopted a less stringent criterion to identify EV cargo proteins that may play critical roles in HNSCC metastasis but not meet the adjusted p-value cutoff. Therefore, we used a log2-fold change criterion of (|normalized log2-fold change| > 1). Under these parameters, 80 and 68 enriched proteins were respectively identified in MOC2-SEVs (log2-fold change >1) and MOC1-SEVs (log2-fold change <-1) (Supplementary Tables 2 and 3).

To analyze the differentially expressed proteins, we conducted gene ontology (GO) analysis on the 80 significantly differentially expressed proteins identified in MOC2-SEVs proteome using STRING and Gene Set Enrichment Analysis (GSEA) (Supplementary Table 2) (25, 26). STRING network analysis of MOC2-SEVs proteome revealed distinct clusters of interacting proteins associated with key biological processes and Reactome pathway analysis, including extracellular matrix (ECM) remodeling, cell migration, neurogenesis/nerve outgrowth, and angiogenesis, all of which are processes associated with tumor metastasis (Figure 1A, Supplementary Table 4). This analysis revealed potential hub and interacting proteins that may act to drive these pathways. Consistent with this analysis, GSEA identified significant enrichment of the hallmark pathways “neuronal system” and “ECM organization” (Figure 1B).

**Figure 1.**
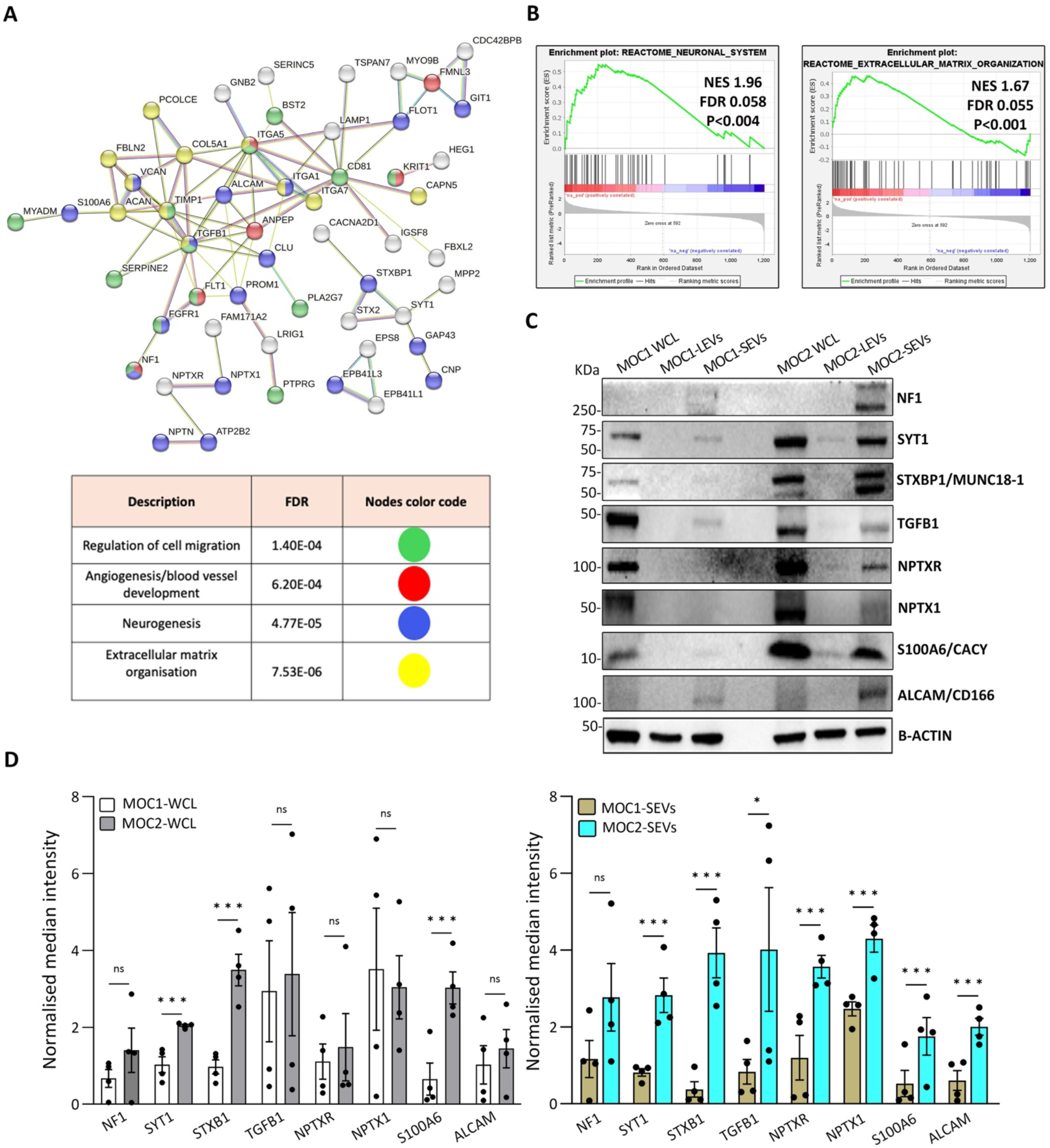
Proteomic profiling of cargoes enriched in MOC2 SEVs. **(A)** A comprehensive protein-protein interaction (PPI) analysis of cargoes enriched in MOC2-derived SEVs was performed using the STRING database to identify and visualize the functional relationships among the proteins within these vesicles. The lower panel represents Gene Ontology (GO) and Reactome pathway categories for significantly enriched proteins, along with their FDR values. Color coding corresponds to different functional categories: red for angiogenesis/blood vessel development, blue for neurogenesis, green for regulation of cell migration, and yellow for extracellular matrix organization, as illustrated in the upper panel. **(B)** Top Reactome pathways were identified using Gene Set Enrichment Analysis (GSEA), based on normalized enrichment score (NES), FDR, and p-values. The NES (green curve) indicates whether a given pathway is significantly enriched at the top or bottom of the ranked protein list. Black vertical lines mark the positions of proteins from each pathway within the ranked list. **(C)** Western blot analysis of whole-cell lysates (WCL), large extracellular vesicles (LEVs), and small extracellular vesicles (SEVs) isolated from MOC1 and MOC2 cells, showing the expression of selected proteins. β-actin was used as a loading control for both cell lysates and EVs. **(D)** Densitometric quantification of immunoblots from at least three independent experiments. Protein levels were normalized to β-actin for SEVs and cell lysates. Data are presented as mean ± SEM. *p <0.05, **p <0.01, ***p <0.001. Statistics, two-tailed unpaired *t* test.

From the upregulated MOC2 EV cargoes, we chose eight to validate by Western blot: neurofibromin 1 (NF1), synaptotagmin-1 (SYT1), syntaxin-binding protein 1 (STXBP1/MUNC18-1), transforming growth factor beta-1 (TGFB1), neuronal pentraxin receptor (NPTXR), neuronal pentraxin-1 (NPTX1), protein S100-A6 (S100A6), and activated leukocyte cell adhesion molecule (ALCAM/CD166). We performed Western blot analysis on both cell lysates and EV samples from MOC1 and MOC2 cells, loading equal protein amounts in each lane and using β-actin as a loading control. Selected proteins predicted to be enriched by the iTRAQ proteomics analysis were indeed significantly enriched in MOC2 EVs compared to MOC1 EVs, with the exception of NF1 (Figure 1C, D). In many cases, the enrichment appears to result from selective EV sorting of those cargoes, as only SYT1, STXBP1, and S100A6 were enriched in MOC2 cell lysates compared with MOC1 lysates (Figure 1C, D).

We also performed STRING and GSEA analysis on the 68 proteins enriched in MOC1-SEVs (Supplementary Table 3). We found that MOC1-SEVs are enriched in protein categories associated with cell adhesion, RNA binding, and pathways linked to infectious disease (Supplementary Figure 2, Supplementary Table 5). Due to the high enrichment of RNA-binding proteins in MOC1-SEVs (Supplementary Figure 2A, B, Supplementary Table 5), we hypothesized that they may carry more RNA than MOC2-SEVs. Indeed, quantitation of the RNA content of purified EVs and cells by absorbance at 260 nm revealed enrichment of RNA in MOC1 EVs compared to MOC2 EVs, whereas the opposite was true for the level of RNA in cells (Supplementary Figure 2C).

We also chose several MOC1 EV-enriched cargoes for Western blot validation: E-cadherin (CDH1), pantetheinase (VNN1), 5’-nucleotidase (CD73/NT5E), 14-3-3 protein sigma (SFN), large ribosomal subunit protein 18 (RPL5), and eukaryotic translation initiation factor 6 (EIF6) (Supplementary Table 4). Of these, CDH1, CD73, SFN, RPL5 and EIF6 were significantly enriched in MOC1-SEVs compared to MOC2-SEVs, however, only CDH1 was significantly enriched in MOC1 cell lysates compared to MOC2 cell lysates, indicating a selective enrichment of CD73, SFN, RPL5, and EIF6 in MOC1 SEVs rather than the parental cells (Supplementary Figure 2D, E).

### Exosome secretion controls primary tumour growth in the murine tongue

To investigate the functional role of SEVs in promoting cancer progression, we generated HRS (hepatocyte growth factor-regulated tyrosine kinase substrate/HGS) knockout MOC2 cell lines using the CRISPR-Cas9 genome editing system (27). HRS is a key component of the ESCRT complex involved in endosomal sorting and exosome/SEV biogenesis (11, 28–30). We confirmed the loss of HRS expression in two independent MOC2 knockout clones by Western blot analysis of cell lysates and purified SEVs (Figure 2A). Classical SEV markers (TSG101, CD63, Flotillin-1) were unaffected by HRS deletion, and β-actin served as a loading control (Figure 2A). TEM and NTA showed no obvious morphological or size distribution differences in SEVs between knockout and control cells, respectively (Figure 2B, C). However, NTA quantification demonstrated a significant reduction in the number of SEVs released from HRS-KO cells relative to controls (Figure 2D), indicating impaired SEV biogenesis despite preserved marker expression and morphology.

**Figure 2.**
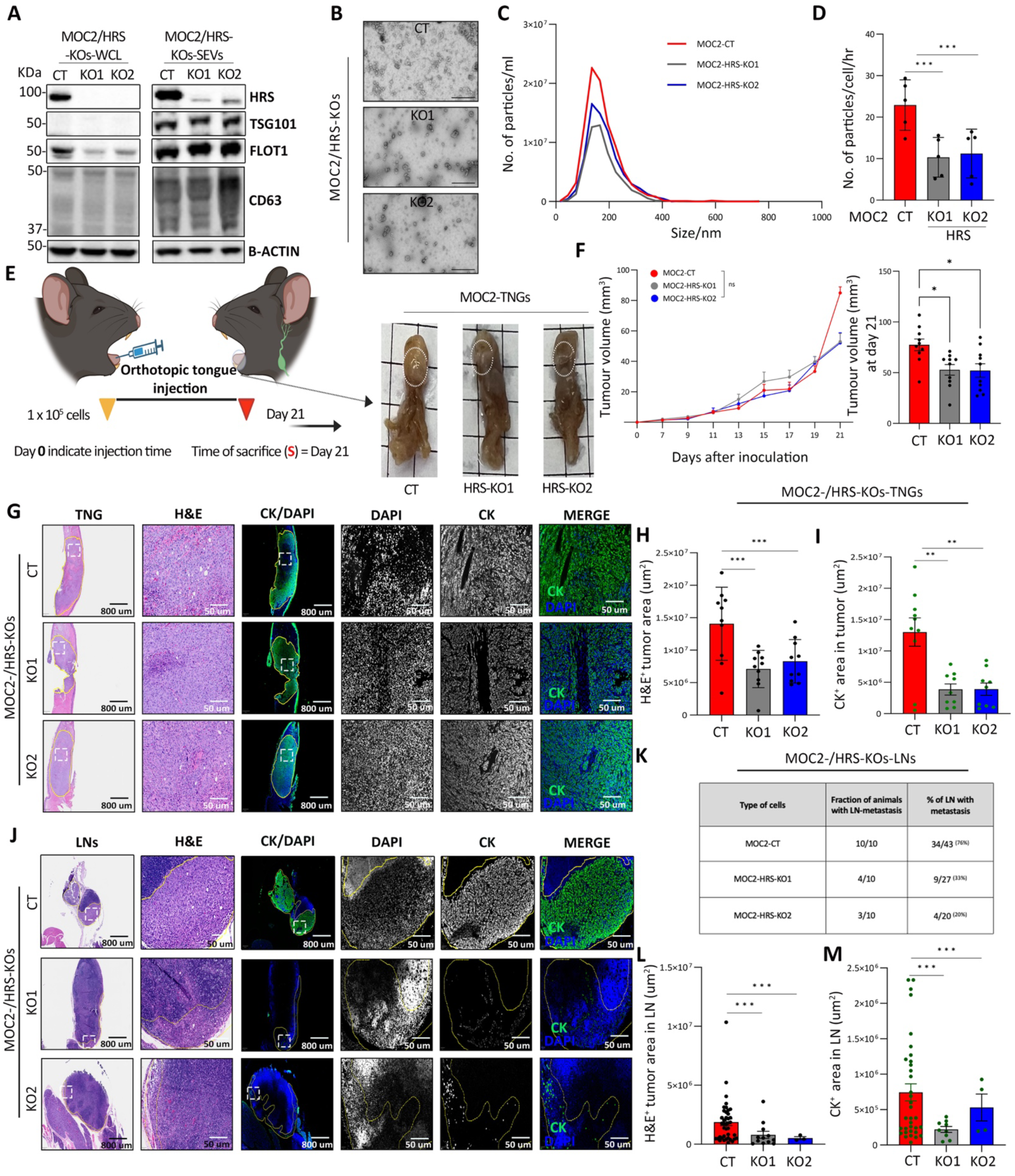
HRS deletion impairs the secretion of SEVs and suppresses lymph node metastasis in oral carcinoma. **(A)** Western blot analysis confirming the successful knockout of the HRS gene in MOC2 cells. Classical SEV markers TSG101, FLOT1, and CD63 were also probed and were enriched in the SEV samples. β-actin served as a loading control. **(B)** Representative transmission electron microscopy (TEM) images of SEVs isolated from MOC2 control (CT) and HRS-knockout (HRS-KO) cells. Scale bars: 100 nm. **(C)** Representative nanoparticle tracking analysis profiles of SEVs from MOC2-CT and HRS-KO cells. **(D)** Quantification of SEV secretion rates derived from NTA, based on EVs collected over 48 hours from MOC2-CT and HRS-KO cells. Cell numbers were counted at the time of media collection (n = 5). **(E)** Left: Illustration of the syngeneic, immune-competent mouse model used for orthotopic tongue injections to induce LN metastasis. Right: Representative images showing primary tongue tumors in mice injected with MOC2-CT or HRS-KO cells. **(F)** *In vivo* tumor growth following orthotopic tongue injections, CT (red, n = 10) compared to HRS-KOs (grey and blue, N = 10). Tumor growth curves represent mean ± SEM. **(G)** Representative immunohistochemistry (IHC) images of H&E (left panel) showing tongue (TNG) tumor area and cytokeratin (CK) staining (right panel) of tongue in CT and HRS-KOs. Yellow line in low power images outlines the tumor whereas the white box indicates the zoomed areas. Scale bars: 50 or 800 μm, as indicated. **(H, I)** Quantification of H&E and CK^+^ tumor areas (N= 10 per group). **(J)** Illustrative IHC images of H&E (left panel) and CK^+^ (right panel) area in lymph nodes (LNs). Scale bars: 50 or 800 μm, as indicated. **(K)** Table shows quantification of the fraction of animals with lymph node metastasis (middle column) or of the number of lymph nodes with detectable cancer metastases (right column) in control (CT) or KO tumors. **(L, M)** Bar graphs depicting H&E-positive tumor area and CK-positive staining in LNs. Data are presented as mean ± SEM: statistics, unpaired *t*-test: *p<0.05, **p<0.01, ***p<0.001. Scale bars, as indicated.

To evaluate the *in vivo* impact of SEV depletion, we orthotopically injected syngeneic mice with control MOC2 cells (MOC2-CT) or two independent HRS knockout clones (MOC2-HRS-KO1 and -KO2) directly into the body of the tongue (Figure 2E). By day 7 or 10 post-injection, all mice developed visible tumors within the tongue tissue. The tumor volume was measured regularly, and the final volume was calculated on day 21 when animals were sacrificed, and tongue tumors were harvested for analysis (Figure 2F). Tumor development was consistently observed in all experimental cohorts. Comparative analysis of longitudinal tumor growth revealed no significant differences in the overall tumor volume between control and HRS-KO groups throughout the study period until day 21 (Figure 2F, left panel), at which time the tumor volume in the control group was significantly greater than that of the HRS-KO group (Figure 2F, right panel).

To further analyze the tumors, we performed immunohistochemical staining of the tumors and surrounding tissues. Consistent with the caliper measurements, analysis of hematoxylin and eosin (H&E)-stained tumor sections demonstrated a significantly larger area of control tumors than in HRS-KO counterparts (Figure 2G, H). To further characterize these tumors, we performed fluorescent immunostaining for pan-cytokeratin (CK), an established marker of epithelial tumor cells. Consistent with the other data, tongues from mice bearing HRS-KO tumors had a reduced cytokeratin-positive tumor area relative to tongues from mice bearing control tumors (Figure 2G, I). Together, these observations indicate that, while depletion of SEVs does not prevent the initiation of tumor formation, over time it influences tumor growth.

### Exosome secretion promotes lymph node metastasis of MOC2 cells

Lymph node metastasis is a well-established predictor of poor prognosis in HNSCC (6). To assess the impact of HRS depletion on cervical lymph node metastasis, we evaluated metastatic burden in cervical lymph nodes using H&E and CK immunostaining (Figure 2J). Strikingly, all mice bearing MOC2-CT tumors (10/10) exhibited lymph node metastases, whereas a markedly lower incidence was observed in HRS-deficient groups; 4 out of 10 mice in the HRS-KO1 tumor group and 3 out of 10 in the HRS-KO2 tumor group developed metastases (Figure 2K). We also quantified the total number of lymph nodes positive for metastatic lesions, revealing a significant reduction in HRS-KO-tumor-bearing mice, with 9 out of 27 nodes positive in KO1 and 4 out of 20 nodes positive in KO2, compared to 33 out of 43 nodes positive in the MOC2-CT-tumor-bearing mice (Figure 2K). Quantitative analysis of the tumor-positive regions within metastatic lymph nodes by H&E and CK-positive staining also revealed a substantial decrease in the size of tumor-positive areas in HRS-KO groups relative to controls (Figure 2L, M).

### Exosome secretion promotes nerve and blood vessel recruitment during primary tumour growth

One of the interesting findings from our proteomic analysis of MOC2-derived SEVs was a significant enrichment of proteins associated with neurogenesis and angiogenesis, suggesting potential roles for SEVs in these processes (Figure 1A). To directly investigate this, we performed IHC on tongue tumors from control and HRS-KO MOC2 cells using neurofilament heavy chain (NF-H, a pan-nerve marker) to assess nerve density and CD31 to visualize blood vessel density. We observed significantly decreased nerve and blood vessel density at the primary tumor and adjacent normal tongue sites for HRS-KO tumor-bearing mice compared to control tumor-bearing mice (Figure 3A-C). These data support a functional role for SEVs in recruiting nerves and blood vessels to the tumor microenvironment.

**Figure 3.**
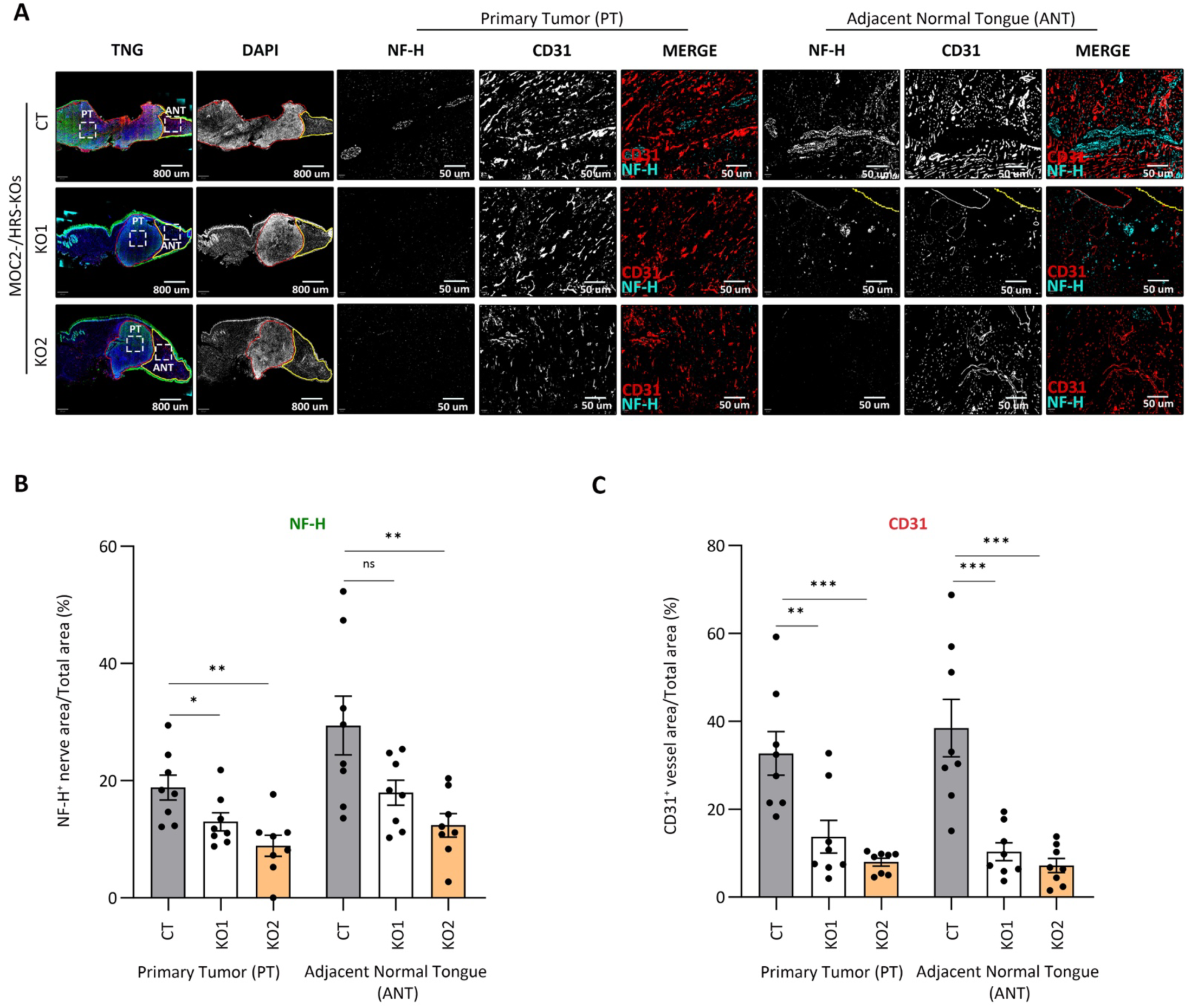
Exosome secretion regulates nerve outgrowth and blood vessel formation within the MOC2 tumor microenvironment. **(A**) Representative immunofluorescence images showing the expression of neurofilament-heavy chain (NF-H), an intermediate filament marker of nerve fibres, and CD31, an endothelial cell marker, within primary tongue tumors. Red demarcation lines outline the primary tumor (PT) region, while yellow lines indicate the boundary separating the adjacent normal tongue (ANT) compartment from the PT. Images highlight the distribution of nerve fibres and vasculature within the TME. **(B, C)** Quantification of NF-H⁺ nerve fibres and CD31⁺ endothelial cells as the percentage of positively stained area in control (CT) and HRS-KO mice (N = 8 mice per group). Data are shown as mean ± SEM, with statistical significance determined using unpaired two-tailed t-tests (*p <0.05, **p <0.01, ***p <0.001). Scale bars: 100 and 800 μm, as indicated.

### Exosome secretion affects the immune microenvironment of MOC2 tumors

Elevated expression of HRS has been observed in tumor tissues and is positively associated with increased levels of circulating SEV-associated PD-L1 in patients with HNSCC (28). In the same study, higher HRS expression was found to inversely correlate with CD8⁺ T cell infiltration in HNSCC tumors, suggesting a role for exosomes in tumor immune evasion (28). Given that the MOC2 murine model of oral squamous cell carcinoma (OSCC) is inherently immunologically “cold,” we tested whether HRS KO modulates tumor immune cell infiltration in our model. Using the immune-specific markers CD8 for cytotoxic T cells and CD45 as a pan-leukocyte marker, we observed a substantial increase in immune cell infiltration within the primary tumor (PT) in HRS-KO tumors compared to control tumors (Figure 4A-C). In contrast, no significant difference in CD8⁺ T and CD45⁺ cell numbers was detected in the surrounding adjacent normal tongue (ANT) regions between HRS-KO and control tumors (Figure 4A-C). These findings are consistent with the literature and suggest that SEVs may promote tumor immune evasion in HNSCC.

**Figure 4.**
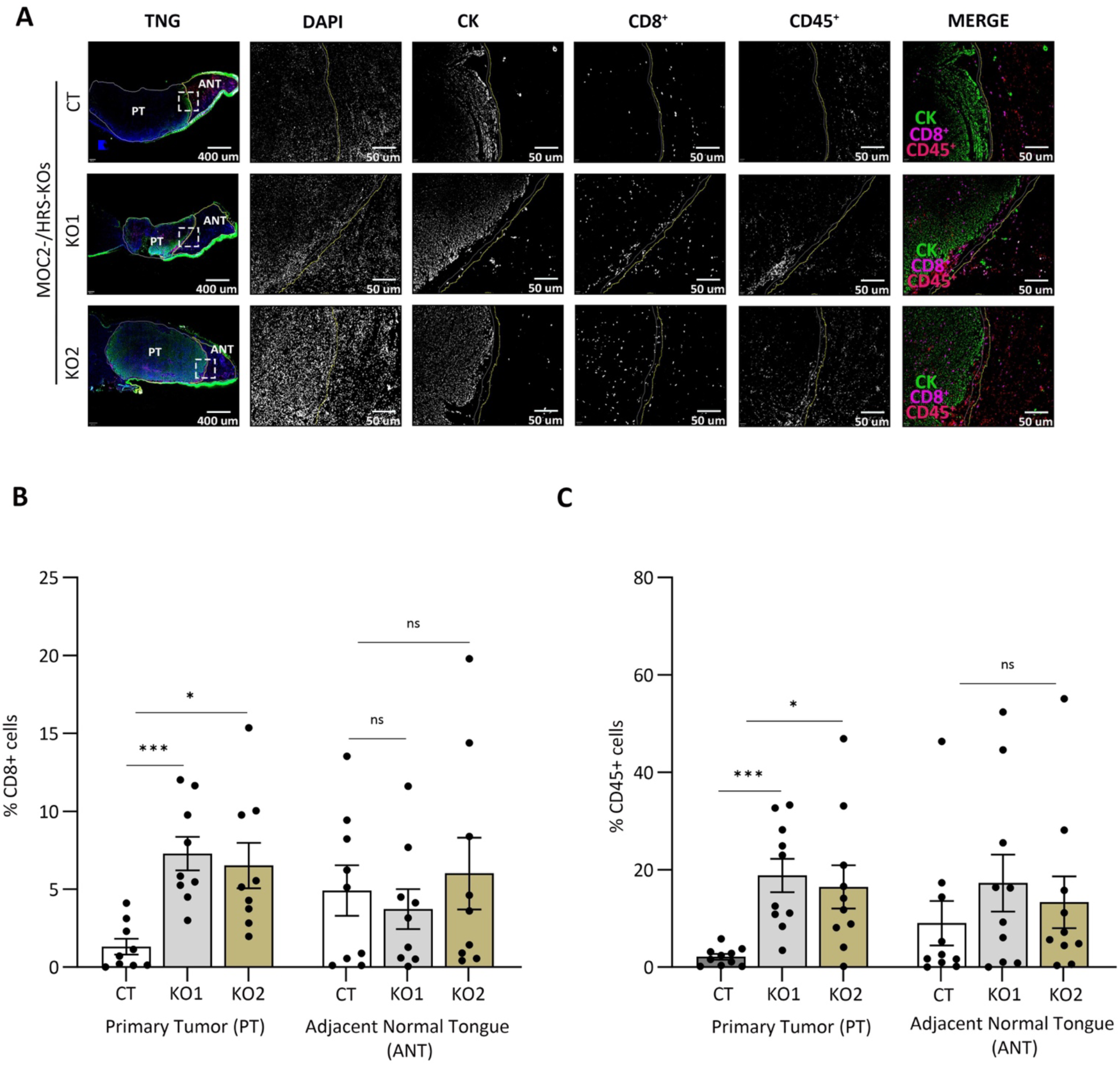
Exosomes regulate immune cell recruitment in the oral tumor microenvironment. **(A)** Representative immunofluorescence images showing the spatial distribution and expression of the epithelial marker cytokeratin (CK), the cytotoxic T cell marker CD8, and the pan-leukocyte marker CD45⁺ within primary tumor (PT) regions and adjacent normal tongue (ANT) compartments. White demarcation lines outline the PT area, while yellow lines indicate the boundary separating the stromal region from the PT. **(B, C)** Quantification of CD8⁺ T cells and CD45⁺ cells as the percentage of positive cells in control (CT) and HRS-KO mice (N = 9 mice per group). Data are shown as mean ± SEM, with statistical analysis performed using unpaired two-tailed t-tests (*p <0.05, **p <0.01, ***p <0.001). Scale bars: 50–400 μm.

### Select protein cargoes enriched in MOC2 SEVs are also elevated in circulating EVs from HNSCC patients and correlate with metastasis

To connect our proteomics and tumor findings with potential prognostic biomarkers based on SEVs, we analyzed gene expression profiles in a cohort of 499 HNSCC patients with RNA-Seq data from the publicly available TCGA database (31, 32). Using Kaplan-Meier analysis for the genes representing each of the 80 enriched proteins in MOC2-SEVs (Supplementary Table 2), we found that overexpression of 9 of the candidate genes in HNSCC tumors was associated with decreased survival of HNSCC patients (Table 1, Figure 5A, Supplementary Figure 3) (33, 34). From those nine proteins, we chose 5 for further analysis: SYT1, STXBP1, TGFB1, ALCAM, and ITGA5 (Figure 5A). These protein candidates were selected based on their statistical association with decreased patient survival and an unfavourable hazard ratio.

**Figure 5.**
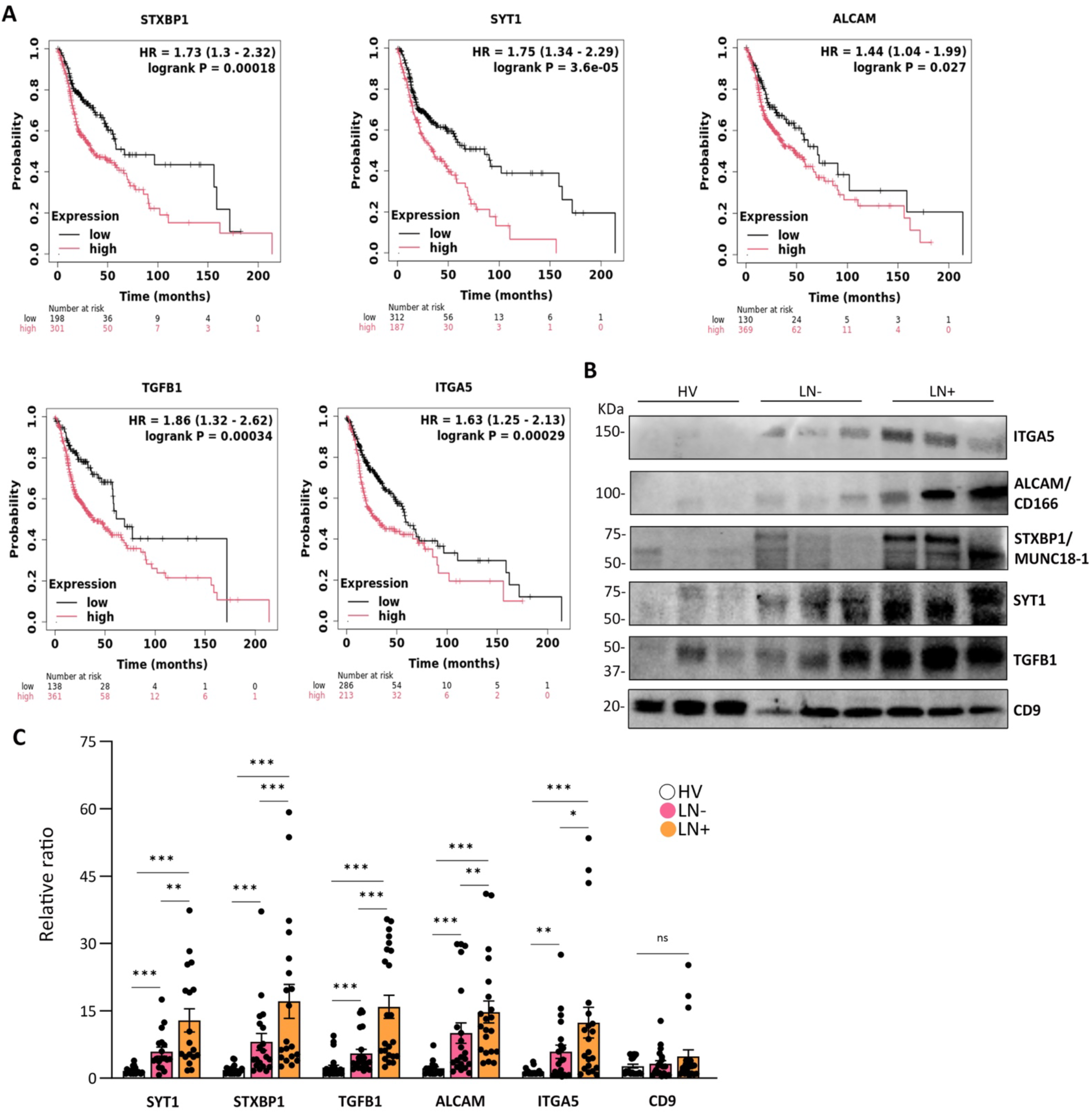
Evaluation of EV cargo expression in SEVs purified from the plasma of HNSCC patients with or without LN metastasis. **(A)** Kaplan-Meier survival curves depicting the overall survival of HNSCC patients stratified into high-and low-expression groups for STXBP1, SYT1, ALCAM, TGFB1, and ITGA5. The number of patients in each group is listed under the plots. HR=Hazard Ratio. Logrank p indicates statistical significance. Data from TCGA RNA Seq for 499 patients. **(B)** Representative Western blot analysis showing the expression of SYT1, STXBP1, TGFB1, ALCAM, and ITGA5 in SEVs derived from healthy volunteers (N =15), LN-negative (LN-), and LN-positive (LN+) HNSCC patient plasma samples (N = 18-23, depending n the blot). **(C)** Quantitative bar graphs show the Western blot densitometric analysis of SYT1, STXBP1, TGFB1, ALCAM, ITGA5, and CD9 levels across the three groups (HV, LN-, LN+). The relative ratio of each protein candidate was determined by quantifying the intensity values obtained from Western blots. For normalization, a single reference intensity value from HV was consistently used across each individual blot. This approach allowed us to standardize protein expression levels and calculate relative ratios in a reproducible manner, ensuring comparability across samples. Data are presented as mean ± SEM, and statistical significance was determined using an unpaired two-tailed *t-test* (*p <0.05; **p <0.01; ***p <0.001).

**Table 1.** Pro-metastatic molecular factors enriched in SEVs released by aggressive MOC2 cells.

| Gene<sup>1</sup><br>name | Description<sup>1</sup> | Hazard<br>Ratio<sup>2</sup> | P-value<sup>2</sup> | Normalized<br>Log2 MOC2-SEVs/MOC1-sEVs<sup>3</sup> |
| --- | --- | --- | --- | --- |
| SYT1 | Synaptotagmin-1 | 1.75 | 0.00036 | 1.9145 |
| TIMP1 | Metalloproteinase inhibitor 1 | 1.53 | 0.0023 | 1.6175 |
| TGFB1 | Transforming growth factor beta-1 | 1.88 | 0.00034 | 1.5085 |
| ITGA5 | Integrin alpha-5 | 1.63 | 0.00029 | 1.4355 |
| NPTN | Neuroplastin/Stromal cell-derived receptor<br>1 (SDR-1/SDFR-1) | 1.52 | 0.0067 | 1.2675 |
| GAP43 | Growth associated protein<br>43/Neuromodulin/pp46 | 1.40 | 0.02 | 1.2605 |
| ALCAM | Activated leukocyte cell adhesion<br>molecule/CD166 antigen | 1.44 | 0.0027 | 1.0835 |
| MYADM | Myeloid-associated differentiation marker | 1.44 | 0.01 | 1.0505 |
| STXBP1 | Syntaxin-binding protein 1/MUNC18-1 | 1.73 | 0.00018 | 1.0035 |

<sup>1</sup>Gene names and protein descriptions were retrieved from the UniProt database. <sup>2</sup>Hazard ratios and P-values were obtained from the Kaplan–Meier Plotter online platform. <sup>3</sup>Log2 fold-change values represent the iTRAQ-based quantitative protein ratio between MOC2-SEVs and MOC1-SEVs (see Table S1). Proteins with $\log_2(\text{MOC2-SEVs/MOC1-SEVs}) \geq 1$ (i.e., $\geq 2$ -fold enrichment) were classified as selectively enriched in MOC2-SEVs compared with MOC1-SEVs.

To determine whether expression of these 5 proteins in circulating SEVs isolated from patient plasma are associated with lymph node metastasis, we analysed deidentified plasma samples from 18-23 patients with HPV-negative metastatic oral carcinoma with (LN+) or without (LN−) known lymph node involvement, and 15 healthy volunteers (Table 2). To avoid major differences due to tumor size, all of the HNSCC samples were chosen from patients with stage III and IV tumors.

**Table 2.**
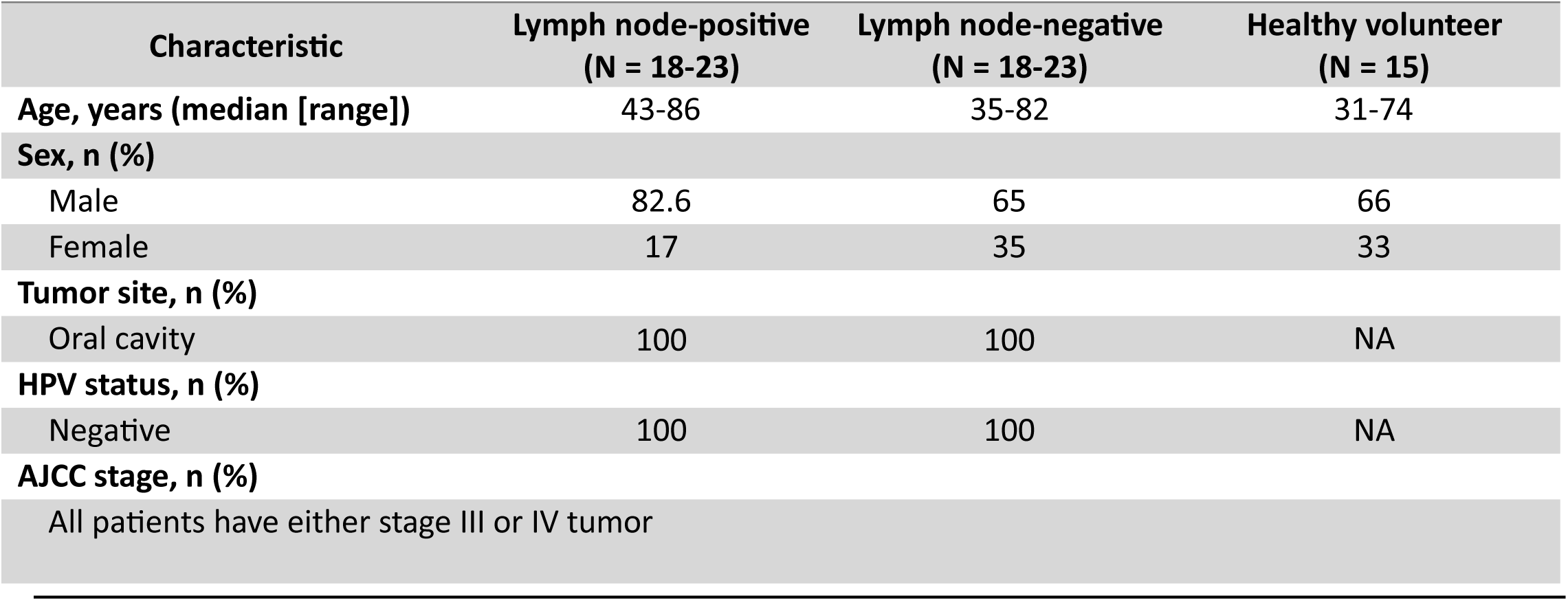
Clinicopathological characteristics of HNSCC patient cohort.

| **Characteristic** | **Lymph node-positive**<br>**(N = 18-23)** | **Lymph node-negative**<br>**(N = 18-23)** | **Healthy volunteer**<br>**(N = 15)** |
| --- | --- | --- | --- |
| **Age, years (median [range])** | 43-86 | 35-82 | 31-74 |
| **Sex, n (%)** |  |  |  |
| Male | 82.6 | 65 | 66 |
| Female | 17 | 35 | 33 |
| **Tumor site, n (%)** |  |  |  |
| Oral cavity | 100 | 100 | NA |
| **HPV status, n (%)** |  |  |  |
| Negative | 100 | 100 | NA |
| **AJCC stage, n (%)** |  |  |  |
| All patients have either stage III or IV tumor |  |  |  |

SEVs were isolated from patient plasma using a commercially available affinity isolation kit targeting the tetraspanins CD9, CD63, and CD81. We then assessed the expression of our selected biomarker candidates in SEVs using Western blot analysis. Quantitative analysis of Western blots revealed that SYT1, STXBP1, TGFB1, ALCAM, and ITGA5 were significantly elevated in SEVs from both lymph node-negative (LN−) and lymph node-positive (LN+) patients compared to healthy volunteer-SEVs. Moreover, SYT1, STXBP1, TGFB1, ALCAM and ITGA5 exhibited significantly higher expression in lymph node-positive (LN+) patient-SEVs relative to lymph node-negative (LN−) patient SEVs (Figure 5B, C). CD9 expression levels in plasma-SEVs were not significantly different among healthy volunteer (HV), lymph node-negative (LN−) and lymph node-positive (LN+) HNSCC patients, indicating that SEVs are consistently present across these groups, but the cargo proteins are enriched in HNSCC SEVs.

## Discussion

In this study, we employed a multipronged approach to identify mechanisms by which SEVs secreted from oral carcinoma cells drive lymph node metastasis. We identified exosome secretion as a driver of lymph node metastasis in an orthotopic syngeneic mouse model of HNSCC (21, 35). We also employed iTRAQ-based proteomics to identify SEV cargoes associated with aggressive HNSCC phenotypes, including induction of angiogenesis and neurogenesis. Finally, we integrated our proteomic findings with analysis of publicly available data to identify a panel of proteins enriched in circulating EVs from HNSCC patients with LN–metastatic disease.

The comparative proteomics analysis of SEVs derived from metastatic versus non-metastatic cell lines identified multiple EV cargoes associated with cancer progression, including angiogenesis, ECM remodeling, and neurogenesis. Nine of these protein candidates correspond to genes that, when overexpressed in HNSCC tumors, are associated with decreased overall survival in the TCGA dataset. Of those nine candidates, we analyzed the levels of five proteins in circulating patient SEVs by Western blotting. All of them were enriched in plasma SEVs from HPV-negative HNSCC patients compared to healthy controls. Furthermore, all five SEV proteins were present at higher levels in SEVs from LN-positive HNSCC patient samples compared to SEVs from LN-negative patient samples. These findings suggest that SEVs might be a useful platform for liquid biopsy to identify patients at risk of lymph node metastasis. Future directions include validating these findings in an independent cohort, carrying out a prospective comparison between surgical lymph node staging and liquid biopsy, and exploring the remaining four candidate biomarkers for their association with HNSCC metastasis.

Many of the candidate proteins that were enriched in circulating HNSCC SEVs have known roles in cancer progression. For example, TGF-β1 is a master regulator of the tumor microenvironment, driving immune suppression, EMT, fibrosis, and therapy resistance (36, 37). Recent reports from the Whiteside group demonstrated elevated levels of TGF-β1 in plasma-derived exosomes from HNSCC patients (38), and its overexpression has also been reported in blood, breast, pancreatic, and colorectal cancers, where it promotes invasion and immune evasion (39, 40). Along with PD-L1 (41, 42) and other immune regulators, it could contribute to the immune exclusion phenotype that we observed to be regulated by exosomes in our orthotopic HNSCC tumor model.

Two other candidate EV cargoes, ALCAM and integrin α5 (ITGA5) are integral to cell adhesion and invasion, facilitating metastatic spread (43–45). ITGA5 is also overexpressed and associated with poor prognosis in glioma, gastrointestinal, and NSCLC (46–48). ALCAM has been implicated in endometrial, pancreatic, and prostate cancer progression and has been suggested as a predictive marker for recurrence (49–51).

The final two EV cargo candidates associated with HNSCC patient metastatic status, SYT1 and STXBP1/MUNC18-1, are SNARE complex proteins that bind each other and are known for their roles in neuronal synaptic vesicle trafficking and release (52–54). By contrast, their roles in cancer cells are much less understood. SYT1 is a vesicle-anchored protein that acts as a primary Ca²⁺ sensor to trigger rapid, Ca²⁺-dependent vesicle fusion (55, 56).

STXBP1/MUNC18-1 acts as a key chaperone and regulatory adaptor for syntaxin-1, orchestrating SNARE complex assembly and membrane fusion dynamics (54). STXBP1 plays an essential role in synaptic vesicle exocytosis and neurotransmitter release, and its disruption has been linked to profound neurological consequences, including impaired synaptic transmission and severe cognitive deficits in patients (57). A recent report indicates that STXBP1 plays an active role in shaping an immunosuppressive TME in HNSCC, through promoting the secretion of multiple chemokines (CXCL1/2/5/8) for immunosuppressive myeloid cells. In addition, STXBP1 facilitates the membrane localization of CD47, a critical ‘don’t eat me’ signal that enables tumor cells to evade phagocytosis by macrophages (58). Together with our findings that SYT1 and STXBP1 are enriched on SEVs from metastatic patient samples and mouse cell lines, and their known roles in vesicle trafficking, we speculate that SYT1 and STXBP1 may be involved in the secretion of EVs and cytokines from HNSCC tumors. A future direction is to test this hypothesis.

To directly test the role of exosomes in HNSCC metastasis, we inhibited exosome biogenesis in MOC2 cells by deleting the HRS gene (17). Loss of HRS had a mild effect on primary tumor growth and a strong effect on LN metastasis. Consistent with our proteomic findings, exosome secretion greatly impacted both blood vessel and nerve density, with decreases in HRS KO tumors. While our proteomics results did not identify many immune regulatory proteins (with the exception of TGF-β1), we also observed enhanced infiltration of CD8⁺ T cells and CD45⁺ immune populations inside HRS-KO tumors, suggesting that tumor-derived SEVs contribute to immune exclusion phenotypes in HNSCC. This matches previous studies in melanoma linking HRS phosphorylation to poor immune infiltration and resistance to immune checkpoint therapy (29). Overall, our results suggest that SEVs are not just passive carriers of tumor biomarkers but also active players in the process of metastasis and immune modulation.

Overall, our findings further implicate small EVs in metastatic cancer spread and suggest that plasma SEV profiling could be a powerful, non-invasive tool to detect LN metastasis in HNSCC. Our approach has the potential to identify new biomarkers of disease, with the ultimate goal of developing SEV-based liquid biopsies that could enable personalized treatment plans while reducing treatment-related toxicity.

## Methods

### Cell lines and culture

The MOC1 and MOC2 cell lines were obtained from Dr. Young Kim (Vanderbilt University) and maintained in a 2:1 mixture of Iscove’s Modified Dulbecco’s Medium (IMDM, Gibco^TM^, cat. # 12440053) and Ham’s F12 (Gibco^TM^, cat. # 11-765-062), supplemented with 10% FBS, 5 ng/mL epidermal growth factor (EGF; biotechne, cat. # 236-EG), 400 ng/mL hydrocortisone, and 5 mg/mL insulin (Sigma-Aldrich, cat. # 11070-73-8). All cells were cultured in T75 flasks at 37°C in 5% CO_2_, passaged every 4 days, and routinely tested negative for mycoplasma contamination. Cell viability was monitored by trypan blue staining. Authentication was confirmed by STR profiling every 6 months.

### Antibodies

Information about antibodies used in this study is provided in Supplementary Table 6.

### EV isolation from conditioned medium

Isolation and analysis of EVs was performed according to MISEV standards (59, 60). Briefly, cells were cultured to ∼80% confluence, washed three times with 1x PBS, and incubated for 48 hours in Opti-MEM (Gibco^TM^, cat. # 11058-021). Conditioned medium was centrifuged sequentially at 300 × *g* for 5 minutes, 2,000 × *g* for 25 minutes, and 10,000 × *g* for 30 minutes to remove cells, debris, and large extracellular vesicles. SEVs were purified from the supernatant by two methods:

1. For proteomics analysis, SEVs were isolated by cushion density gradient (22). Briefly, the supernatant was overlaid onto a 2-mL 60% iodixanol cushion and subjected to ultracentrifugation at 100,000 × *g* for 18 h using an SW32 rotor (Beckman Coulter). Following centrifugation, the bottom 3 mL comprising the 1 mL EV-containing fraction mixed with 2 mL iodixanol (final concentration 40%) was transferred to the bottom of a new ultracentrifuge tube. Sequential layers of 20%, 10%, and 5% iodixanol were then carefully overlaid. Iodixanol solutions were prepared by diluting OptiPrep (60% aqueous iodixanol) with 0.25 M sucrose and 10 mM Tris (pH 7.5). The gradient was centrifuged for an additional 18 h at 100,000 × *g*, after which 12 density fractions were collected. Fractions were diluted in PBS and pelleted by ultracentrifugation at 100,000 × *g* for 3 h. EVs from fractions 6 and 7 were pooled and used for downstream analyses as SEVs (24).
2. For all other experiments from conditioned media, including for Western Blot analysis, NTA and TEM, crude small EVs (SEVs) were isolated from supernatants by ultracentrifugation at 100,000 × *g* for 3 hours in a Ti45 rotor (Beckman Coulter, cat. # 339160), then further purified by subjecting the pellet to flotation density gradient ultracentrifugation. Briefly, pelleted SEVs were resuspended in 40% wt/vol iodixanol (OptiPrep, supplier, cat. #), transferred to a new tube, and overlaid with 5%, 10%, 20%, and 40% wt/vol iodixanol in 0.25 M sucrose/10 mM Tris, pH 7.5. Gradients were centrifuged at 100,000 × *g* for 18 hours in a SW40 rotor (Beckman Coulter, cat. # 331302). Twelve fractions were collected, diluted in PBS, pelleted again at 100,000 × *g* for 3 hours in a TLA 110 rotor (Beckman Coulter, cat. #366735) and resuspended in PBS. SEV-enriched 6 and 7 fractions (1.10–1.18 g/mL) were combined and probed for classical EV marker enrichment (TSG101, CD9, CD63, Flotillin) (61).

### SEV isolation from human plasma

Plasma samples (1 mL aliquots) were obtained from HNSCC Biorepository at Vanderbilt University Medical Center under IRB protocol ##030062 and stored at −80°C. SEVs were isolated using the EasySep Human Pan-Extracellular Vesicle Positive Selection Kit (STEMCELL Technologies, cat. # 17891) according to the manufacturer’s protocol. To enhance the purity of SEVs, we modified the manufacturer’s protocol by incorporating additional washing steps and increased incubation time aimed at depleting excess plasma albumin, which is highly abundant in plasma. First, each plasma sample was diluted 1:1 in DPBS. Plasma samples were first clarified by centrifugation at 10,000 × g for 30 min, after which the supernatant was transferred to a 5-mL polystyrene tube. A 50 μL antibody cocktail was added, mixed thoroughly, and incubated for 30 min, followed by the addition of 100 μL RapidSpheres and a further 30-min incubation. Samples were then brought to a final volume of 2.5 mL by adding 1.5 mL buffer and gently mixed by pipetting. Tubes were placed uncapped into a magnetic rack and incubated for 20 min, after which the supernatant was carefully discarded unless retained for downstream analyses. The bead-bound EVs were washed four times by resuspension in 2.5 mL buffer, incubated for 3 min on the magnet, and removal of the supernatant demonstrateed at each step. Finally, tubes were removed from the magnet and gently flicked to release EVs from the tube wall into the remaining sample volume at the bottom for downstream processing.

### Size distribution analysis of EVs

Particle size distribution and concentration were assessed as previously described (24) using a ZetaView® PMX 110 NTA instrument (Particle Metrix GmbH). Calibration was performed with 100 nm polystyrene beads prior to each session. Samples were diluted 1:500 in PBS, analyzed in triplicate at 25°C, and recorded using identical, standardized acquisition parameters under controlled temperature conditions (25°C).

### Morphological analysis of EVs

Formvar carbon-coated copper grids (FCF-200-Cu) were sequentially washed with Milli-Q water and 100% ethanol, with excess liquid carefully removed by filter paper wicking between steps. Ten microliters of purified EVs (1 µg/µL) were adsorbed onto grids for 30 minutes, negative stained with 2% phosphotungstic acid (pH 6.1) for 20 seconds and air-dried overnight. Images were acquired at 110,000× magnification using a FEI Tecnai T12 TEM equipped with a 120 kV LaB6 source, Gatan cryo-transfer stage, and AMT XR41-S CCD camera (2K × 2K resolution), and representative fields were analyzed for vesicle morphology and integrity (62).

### Western blot analysis and protein concentration measurement

For sample preparation, EVs and cell lysates were solubilized in SDS sample buffer (4% SDS, 20% glycerol, 0.01% bromophenol blue, 0.125 M Tris-HCl, pH 6.8) containing 100 mM DTT as reducing agent. Protein quantification was achieved through 1D SDS-PAGE followed by SYPRO Ruby staining and densitometric analysis using an iBright image Systems (19). Protein samples (20 μg) were separated by electrophoresis using 4-12% Bis-Tris NuPAGE gels (Thermo Fisher) with MES running buffer at 150 V for 1 hour. Following separation, proteins were transferred to nitrocellulose membranes using the iBlot Dry Blotting System (Life Technologies) at 12 V for 10 minutes. Prior to immunodetection, membranes were washed three times (15 min each) with TBST buffer. Protein detection was performed using enhanced chemiluminescence (Thermo Scientific, cat. # 32106) with 5-minute incubation followed by imaging on an iBright imaging system (ThermoFisher Scientific).

### RNA extraction

Total RNA was isolated from equal numbers of small extracellular vesicles (SEVs) and an equal number of matched parent cells using the miRNeasy Mini Kit (Qiagen, Valencia, CA, USA; Cat# 217004), following the manufacturer’s protocol under RNase-free conditions. For SEV samples, RNA extraction was performed immediately following vesicle quantification to prevent degradation. Parentals cells were harvested in parallel to ensure comparable physiological and culture conditions.

Briefly, <u>S</u>EV or cell pellets were lysed in QIAzol lysis reagent with vigorous mixing, followed by phase separation with chloroform. The aqueous phase was transferred to RNeasy spin columns, washed, and eluted in RNase-free water. The miRNeasy chemistry enables efficient isolation of small RNAs <200 nt, including microRNAs and other regulatory small RNAs, as well as total RNA.

RNA concentration and purity were measured using a NanoDrop 2000 spectrophotometer (Thermo Fisher Scientific) by recording absorbance at 260 nm (A260). Total RNA amount (ng) was calculated as the product of the measured concentration (ng/µl) and the elution volume. To normalize RNA content to particle or cell number, total RNA (ng) was divided by the total number of SEVs (quantified by nanoparticle tracking analysis) or the total number of cells (determined by automated cell counting), yielding RNA per EV (ng) or RNA per cell (ng), respectively.

All RNA samples were stored at −80 °C until further analysis.

### Mass spectrometry-based protein identification and quantification

Lysis of EVs, protein digestion, iTRAQ labeling, and quantitative LC-MS/MS analysis: MOC1 and MOC2-SEVs were solubilized in 100 mM TEAB, 300 mM NaCl, 2% NP-40, 0.5% Deoxycholate. Samples were sonicated twice for 10 min each in ice-cold water using an ultrasonic bath (model 97043-988, VWR). Cleared lysates were obtained by centrifugation at 13,500 rpm for 10 min in a Prism R microcentrifuge (Labnet). Protein concentrations were determined with the Pierce BCA protein assay (Thermo Scientific, cat. # 23225).

Proteins were precipitated overnight at −20 °C with ice-cold acetone, centrifuged at 14,000 × g for 30 min at 4 °C, and acetone was removed. Protein pellets were washed with cold acetone and centrifuged, acetone was removed, and proteins were resuspended in 25 ul of 4 M urea in 250 mM TEAB (pH 8.0). Reduction was carried out by addition of 2ul of 50mM TCEP, and cysteine residues were alkylated by addition of 1 µl of 200 mM MMTS. Samples were diluted to 40 uL with 250 mM TEAB and digested overnight with 2ug of sequencing-grade trypsin.

Peptides (25ug) were labeled with iTRAQ reagents (SCIEX) according to the manufacturer’s instructions, using a 1-unit vial of reagent for each sample. iTRAQ reagents were reconstituted in ethanol (90%), and labeling was performed for 2 hours. MOC1 and MOC2 EVs were labeled with iTRAQ reagents 114 and 116. Labeled peptides were combined and desalted using a modified Stage-tip for clean-up. First, a C18 membrane disc (ChromTech C18 SPE Empore disk) was cored with a 16-gauge needle and inserted into a 200 µL pipette tip. C18 resin (Phenomenex Jupiter C18, 5 µm) suspended in methanol was added to the tip, packed by centrifugation, and equilibrated with 0.1% TFA. Peptides were diluted, acidified, and loaded onto the tips by centrifugation. After the tips were washed with 0.1% TFA, peptides were eluted with 80% acetonitrile containing 0.1% TFA. Eluates were dried by speed-vac centrifugation.

Peptides were reconstituted in 0.1% formic acid and loaded onto a self-packed biphasic C18/SCX MudPIT column using a helium-pressurized cell. The MudPIT column (360 o.d. × 150 μm i.d. fused silica) contained 6 cm Luna SCX (5 μm, 100 Å) followed by 4 cm Jupiter C18 (5 μm, 300 Å). After sample loading, the column was coupled via an M-520 microfilter union to a C18 fused silica analytical column (100 μm i.d.) equipped with a laser-pulled emitter tip. The analytical column was packed with 20 cm Jupiter C18 (3 μm, 300 Å) material. Peptides were separated on a Dionex Ultimate 3000 nanoLC using a 13-step salt pulse gradient (0mM, 25mM, 50mM, 75mM, 100mM, 150mM, 200mM, 250mM, 300mM, 500mM, 750mM, 1M, and 2M ammonium acetate), followed by reverse phase chromatography (350 nl/min) using a 95-minute gradient. The mobile phase solvents consisted of 0.1% formic acid/99.9% water (solvent A) and 0.1% formic acid/99.9% acetonitrile (solvent B). For the first 11 SCX fractions, the reverse phase gradient consisted of 2–50 % B for 83 min, 50 % B for 1 min, 50-2 % B for 0.5 min, and 2 % B for 10.5 min (column equilibration). For the last two fractions, the gradient was adjusted to 2–98 % B for 83 min, followed by 98 % B for 1 min, 98-2 % B for 0.5 min, and 2 % B for 10.5 min. Peptides were analyzed using a data-dependent (top 15) method on a Q Exactive Plus Orbitrap mass spectrometer. MS1 spectra were acquired at 70,000 resolution with an AGC target of 3 × 10^6. HCD MS/MS spectra were acquired with 17,500 resolution, an AGC target of 1 × 10^5, and normalized collision energy was set to 30 nce. Dynamic exclusion was enabled using a 30s exclusion duration.

Spectra were searched in Spectrum Mill (version B.04.00, Agilent) against the *Mus musculus* UniprotKB protein database. Search parameters included trypsin specificity, ±20 ppm precursor/product mass tolerance, fixed modifications of cysteine alkylation and iTRAQ labelling (lysine and N-terminus), and variable modification of methionine oxidation. Database search results were filtered to <1% FDR at the peptide and protein level. The median log_2_ iTRAQ ratio for each protein was calculated from all quantified peptides. For statistical analysis, log₂ ratios were first fit to a normal distribution using least-squares regression. The mean of the Gaussian fit was used to normalize individual log2 protein ratios. The mean and standard deviation derived from the Gaussian fit of normalized log2 ratios were then used to calculate *p* values using Z score statistics. Multiple testing correction was performed using the Benjamini–Hochberg procedure (63).

### Database search and quantification

MS/MS spectra were compared against the human subset of the UniProt KB protein database, and Spectrum Mill’s autovalidation features were applied to ensure a false discovery rate (FDR) below 1% at both the protein and peptide levels. For each protein, the median log_2_ iTRAQ ratio was calculated from all identified peptides, and frequency distributions were plotted in GraphPad Prism 6.01. These log₂ ratios, which generally follow a normal distribution, were fitted using least-squares regression. The resulting Gaussian fit provided the mean and standard deviation values, which were then used in Z-score calculations (𝑧 = (𝑥−𝑚)/𝑠) to convert the dataset into a standard normal variable. Using the known properties of the standard normal curve, p-values were derived for each protein ratio. Finally, multiple testing correction was performed using the Benjamini–Hochberg procedure. We performed differential expression analysis on 1257 proteins, which were quantified in iTRAQ experiment. Requiring a normalized Log2 ± 1-fold change, we identified 86 up-regulated proteins and 68 down-regulated proteins.

### Protein-protein interaction and Gene Set Enrichment Analysis (GSEA)

For Gene Ontology (GO) enrichment analysis, the STRING database (version 11.5) (64) was utilized to predict potential protein–protein interactions (PPIs) for differentially expressed proteins. Interaction predictions in STRING are based on multiple evidence sources, including experimental data, computational predictions, and public text collections. A high confidence combined score threshold of > 0.7 was applied to ensure reliable associations. Proteins that did not display any connections to other nodes in the PPI network were excluded from the final visualization to improve interpretability. In addition, Gene Set Enrichment Analysis (GSEA) (https://www.gsea-msigdb.org/) was performed using a web-based platform that offers a variety of predefined gene sets, statistical analysis options, and visualization features. This tool enables the identification of significantly enriched biological pathways, molecular functions, or cellular processes associated with the experimental data (65).

### Mice

Sex was not evaluated as a biological variable in this study, as only female animals were included. Female C57BL/6 mice (6-8 weeks old; Charles River Laboratories) were used for all experiments. Use of a single sex ensured compatible group housing, minimized animal numbers, and maintained experimental consistency in accordance with the 3Rs. Mice were maintained under specific pathogen-free conditions at Vanderbilt University Medical Center (VUMC) in a facility accredited by the American Association for the Accreditation of Laboratory Animal Care (AAALAC), in full compliance with U.S. federal guidelines (USDA, NIH, and PHS). Mice were provided a standard diet of irradiated standard chow (LabDiet) and autoclaved, reverse osmosis–purified water. Environmental conditions (20–25°C, 30–70% humidity, 12-hr light/dark cycles) were controlled to ensure stability in immune and metabolic parameters relevant to tumor progression studies. To minimize pain and distress, animals were anesthetized with isoflurane as needed during tumor volume and body weight measurements. All procedures were approved by the Vanderbilt University Institutional Animal Care and Use Committee (protocol #1800155) and adhered to ARRIVE guidelines.

### Orthotopic HNSCC tumor model establishment

Seven-week-old female C57BL/6 syngeneic mice were used to establish HNSCC tumor models. To generate orthotopic oral tongue tumors, approximately 1 × 10⁵ MOC2 control, MOC2-HRS-KO1, or MOC2-HRS-KO2 cells were trypsinized, washed three times with PBS, and resuspended in 30 µL of serum-free DMEM. Mice were anesthetized with isoflurane, and cell suspensions were then injected directly into the anterior tongues of the mice away from the apex of tongue, and ClearH_2_0 DietGel 76A dietary supplement (Clear H_2_O, cat. # 72-07-5022X) was added in their diet, as previously described (14). Tumor growth and body weight were monitored twice weekly. Tumor size was measured using micro-calipers, and tumor volume was calculated using the formula (A)(B²) π/6, where A represents the longest dimension and B is the perpendicular width. Animal health was carefully maintained by monitoring body weight and hydration status, with supplemental DietGel food provided to minimize potential confounding effects from reduced food or water intake, thereby ensuring that observed outcomes were due to tumor biology rather than systemic health variations. At 21 days post-tumor cell injection, mice were humanely euthanized in accordance with the institutional Animal Care and Use Program (ACUP) guidelines of Vanderbilt University Medical Center. Euthanasia was performed using gradual carbon dioxide (CO₂) inhalation until loss of consciousness, followed by secondary cervical dislocation to ensure death. This two-step method was conducted to minimize animal distress and confirm complete euthanasia prior to tissue collection.

### Immunohistochemistry and Image Analysis

On day 21, tongues and cervical LN were harvested and fixed with 10% neutral buffered formalin solution for 48 hours on a shaker at room temperature. Tissues were then washed in PBS and embedded in paraffin for sectioning by the Translational Pathology Shared Research (TPSR) Core at VUMC. All tissues were sectioned at 5 μm. After serial sectioning, H&E was performed on each tissue, and IHC was performed. Briefly, slides were deparaffinized in xylene and rehydrated in serial ethanol dilutions. Antigen retrieval was performed by heating slides for 17 minutes in Tris EDTA buffer, pH 9, in a pressure cooker at 110°C. Slides were cooled and blocked with 2.5% horse serum (Vector Laboratories). After blocking, slides were incubated overnight at 4°C with a primary antibody in horse serum. Slides were then incubated in anti-rabbit/anti-mouse HRP secondary (Vector Laboratories) for 1hr at room temperature the following day and subsequently incubated in 1:500 Opal 520 (Akoya FP1487001KT) or Opal 570 (Akoya FP1488001KT) for 10 minutes. For serial staining, slides were stripped using Citric Acid buffer, pH 6.1, in a pressure cooker at 110 °C for 2min and then staining was repeated using different antibodies and Opal fluorophores. Slides were mounted using an antifade gold mount with DAPI (Invitrogen). Stained images were acquired using an Olympus SLIDEVIEW VS200 at 20x magnification equipped with an integrated color camera and a high-end Hamamatsu monochrome sCMOS camder with 20x-40x objectives. Images were analyzed with QuPath (version 0.5.1) and quantified using ImageJ software (66).

Briefly, multiplex immunofluorescence images were analyzed using QuPath. Whole-slide images were imported into QuPath, and regions of interest (ROIs) were manually annotated. In ImageJ, individual Opal fluorophore channels were separated and converted to 8-bit grayscale images. Background fluorescence was subtracted, followed by maintaining thresholding across all samples to minimize batch-to-batch variation. For fluorescence quantification, ImageJ measurement parameters were first configured via Analyze-Set Measurements to include Area & Area Fraction. After applying threshold values for each Opal channel, the Analyze-Measure function was used to quantify the area and percentage of area (% area) of positive signal for each ROI. The resulting area and % area-values were exported and used for statistical analysis and generation of corresponding bar graphs.

For cell segmentation and phenotyping, QuPath’s built-in cell detection workflow was applied using default parameters. Briefly, nuclei were identified using a DAPI-based thresholding algorithm, which detects nuclear objects based on intensity and size constraints. Detected nuclei were then expanded by a fixed radius to approximate whole-cell boundaries, allowing assignment of cytoplasmic and/or membrane compartments. Fluorescence intensities for each Opal channel were quantified on a per-cell basis within the defined cellular compartments.

Immune cell populations were identified based on marker-specific fluorescence intensity thresholds, and quantitative metrics including mean fluorescence intensity, cell density, and percentage of marker-positive cells were extracted for downstream statistical analyses. All segmentation and quantification steps were performed using consistent parameters across samples to ensure comparability.

### The Cancer Genome Atlas (TCGA) data analysis and generation of Kaplan-Meier plots

For this study, we used publicly available sequencing data from the HNSC cohort of the TCGA-project (https://portal.gdc.cancer.gov/). A total of 499 primary HNSCC tumor samples with complete and evaluable overall survival data (event status and follow-up time) were included in the final analysis. Of the 528 HNSCC cases available through the Kaplan-Meier Plotter platform, 29 cases were automatically excluded due to missing or incomplete survival annotations. The Kaplan-Meier Plotter integrates gene expression and clinical outcome data derived primarily from The Cancer Genome Atlas (TCGA) Head and Neck Squamous Cell Carcinoma (HNSCC) cohort and applies internal quality control and filtering procedures to retain only samples suitable for time-to-event analysis, thereby ensuring adherence to non-informative censoring assumptions (33, 34, 67).

### Statistics

Statistical tests and the number of replicates for all experiments are indicated in the figure legends. All quantitative data represent at least three independent biological replicates, and error bars represent the standard error of the mean (37), and statistical significance was defined as *P* < 0.05. For comparisons between two independent groups, statistical significance was assessed using a two-tailed unpaired Student’s t-test. For analyses involving more than two groups, differences were evaluated using a one-way analysis of variance (ANOVA), with post-hoc testing as denoted where applicable. For the survival analysis assessing the prognostic significance of the gene of interest, patient samples were stratified into two groups based on median quantile expression levels of the proposed biomarker. Kaplan-Meier survival curves were generated using Kaplan-Meier Plotter online platform to compare overall survival between the high- and low-expression cohorts. The hazard ratio, along with its 95% confidence interval, and the log-rank P < 0.05 were calculated to determine the statistical significance of differences in survival outcomes (33, 34). All analyses were performed using GraphPad Prism 10 (GraphPad Software, Inc.).

### Study Approval

Human plasma samples from deidentified HNSCC patients and healthy volunteers were obtained from the head and neck cancer biorepository at Vanderbilt University Medical Center, operating under IRB-approved protocol #030062. All animal studies were approved by the Vanderbilt University Institutional Animal Care and Use Committee (IACUC) approved protocol #M1800027-02. All animal procedures were conducted in accordance with guidelines approved by the American Association for the Accreditation of Laboratory Animal Care (AAALAC) and were fully compliant with current regulations and standards set by the U.S. Department of Agriculture, the U.S. Department of Health and Human Services, and the National Institutes of Health (NIH).

## Supporting information

Total proteins identified

Highly enriched in MOC2-SEVs

Enriched in MOC1-SEVs

Gene Ontology

Gene Ontology

Antibodies list

## Data Availability

The raw and processed proteomics data have been attached as supplementary table 1 and 2 with all values indicated by different columns. Values for all data points in graphs are reported in the Supporting Data Values file.

## Author contributions

AS and AMW designed the experiments. SS prepared samples for proteomics and AS analyzed the proteomics dataset. BB conducted RNA isolation and analysis. ENA assisted in IHC staining. BB supplied clinical materials and information. AS and AMW wrote the manuscript with input and editing from all coauthors.

## Acknowledgements

This work was supported in part by NIH grant R01-CA249684. We sincerely thank Celine Copeland from the DelGiorno Laboratory for her valuable assistance in scanning and digitizing the immunohistochemistry (IHC) slides, which greatly facilitated image analysis and data visualization. We also extend our gratitude to Dr. Sharon Yang and Miranda K Wilkes from the Translational Pathology Shared Resource (TPSR) Core at Vanderbilt University Medical Center for their technical support, expert guidance, and insightful feedback throughout the histological processing and imaging stages. Their contributions were instrumental in ensuring the rigor and quality of the pathological assessments in this study. We also thank Evan Krystofiak for his support for the TEM imaging of EVs and the Vanderbilt Head and Neck Biorepository for providing plasma samples and deidentified clinical information.

Address correspondence to: Alissa Weaver, Department of Cell and Developmental Biology, Center for Extracellular Vesicle Research, Vanderbilt University School of Medicine, 465 21^st^ Avenue South, Vanderbilt University, PMB 407935 Room T-2212 MCN, Nashville, TN 37240.

## Conflict of interest

The authors have declared that no conflict of interest exists.

---

**Supplementary Figure 1.**
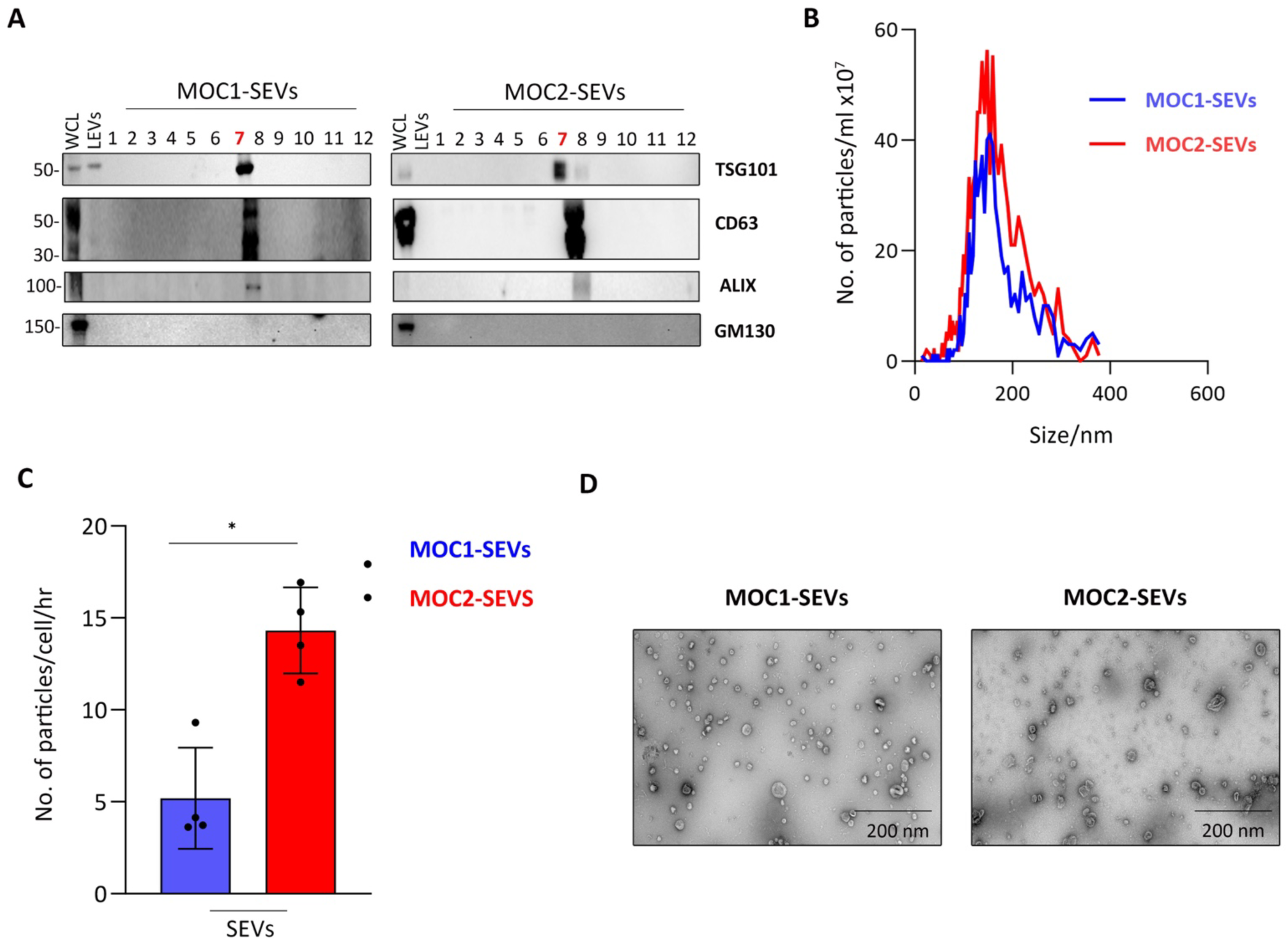
Characterisation of SEVs purified from MOC1 and MOC2 cells. **(A)** Western blot analysis of cell lysates and SEVs fractionated by OptiPrep density gradient ultracentrifugation, confirming enrichment of classical EV markers (TSG101 and CD63) and absence of the negative control marker GM130 in the expected gradient fraction 7. **(B)** Representative nanoparticle tracking analysis profiles illustrating the size distribution and particle concentration of SEVs obtained from MOC1 and MOC2 cell cultures. **(C)** Quantitative comparison of SEV secretion rates, calculated from NTA data, based on vesicles collected over 48 hours from MOC1 and MOC2 cells. Cell counts were obtained at the time of conditioned media collection (n = 3). **(D)** Representative transmission electron microscopy (TEM) images showing the morphology of SEVs isolated from MOC1 and MOC2 cells. Scale bar: 200 nm.

**Supplementary Figure 2.**
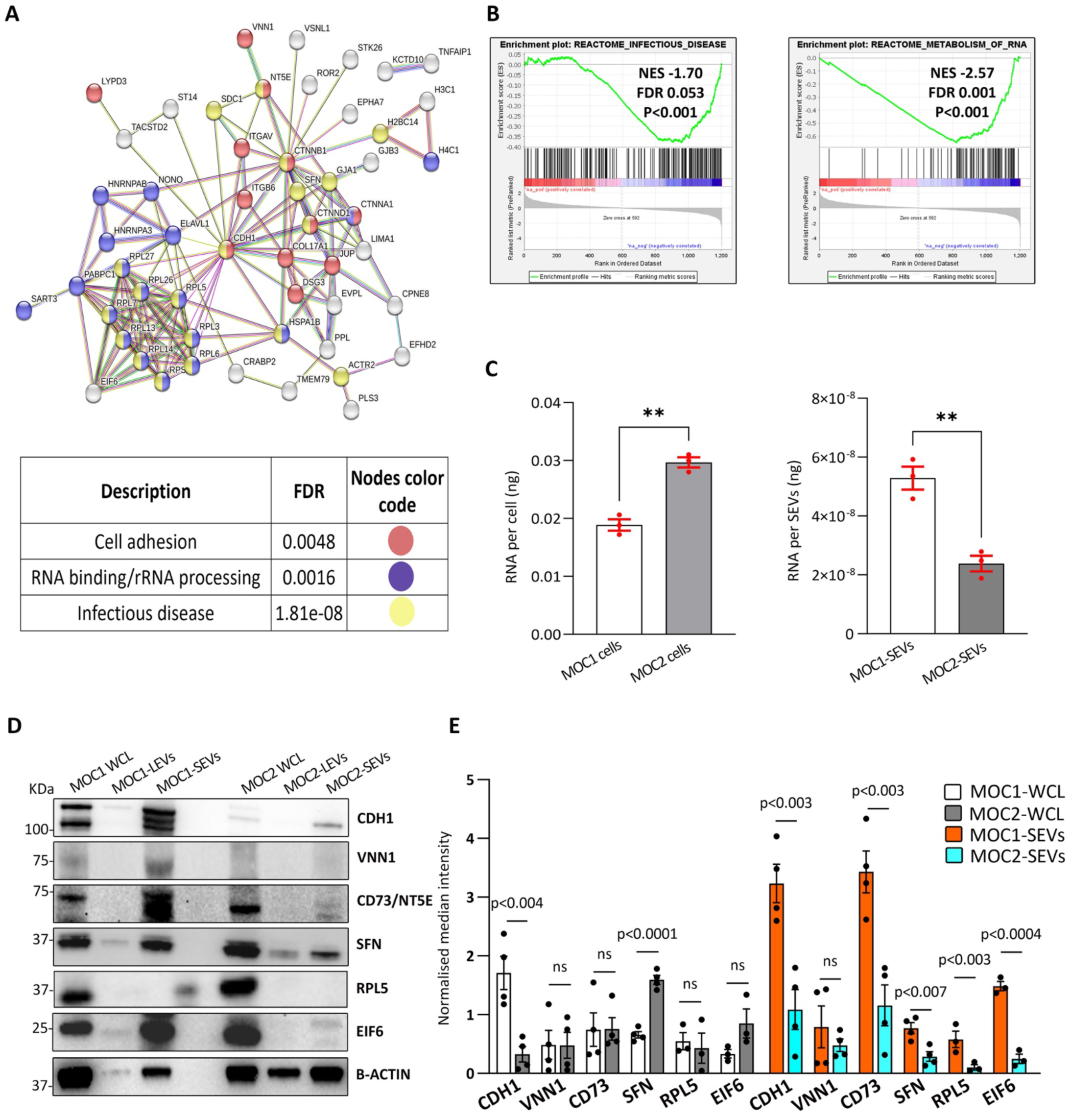
Proteomic profiling of cargoes enriched in MOC1 SEVs. **(A)** A comprehensive protein-protein interaction (PPI) analysis of cargoes enriched in MOC1-derived SEVs was performed using the STRING database to identify and visualize the functional relationships among the proteins within these vesicles. The lower panel represents Gene Ontology (GO) and Reactome pathway categories for significantly enriched proteins, along with their FDR values. Color coding corresponds to different functional categories: red for cell adhesion, blue for RNA binding/rRNA processing, and yellow infectious disease, as illustrated in the upper panel. **(B)** Top Reactome pathways were identified using Gene Set Enrichment Analysis (GSEA), based on normalized enrichment score (NES), FDR, and p-values. The NES (green curve) indicates whether a given pathway is significantly enriched at the top or bottom of the ranked protein list. Black vertical lines mark the positions of proteins from each pathway within the ranked list. **(C)** Bar graphs represent total RNA content, as measured by A260, between MOC1 and MOC2 cells (left panel) and their corresponding SEVs (right panel), (N = 3). **(D)** Western blot analysis of whole-cell lysates (WCL), large extracellular vesicles (LEVs), and small extracellular vesicles (SEVs) isolated from MOC1 and MOC2 cells, showing the expression of selected proteins. β-actin was used as a loading control for both cell lysates and EVs. **(E)** Densitometric quantification of immunoblots from ≥ 3 independent experiments. Protein levels were normalized to β-actin for SEVs and cell lysates. Data are presented as mean ± SEM. *p <0.05, **p <0.01, ***p <0.001. Statistics, two-tailed unpaired *t*-test.

**Supplementary Figure S3.**
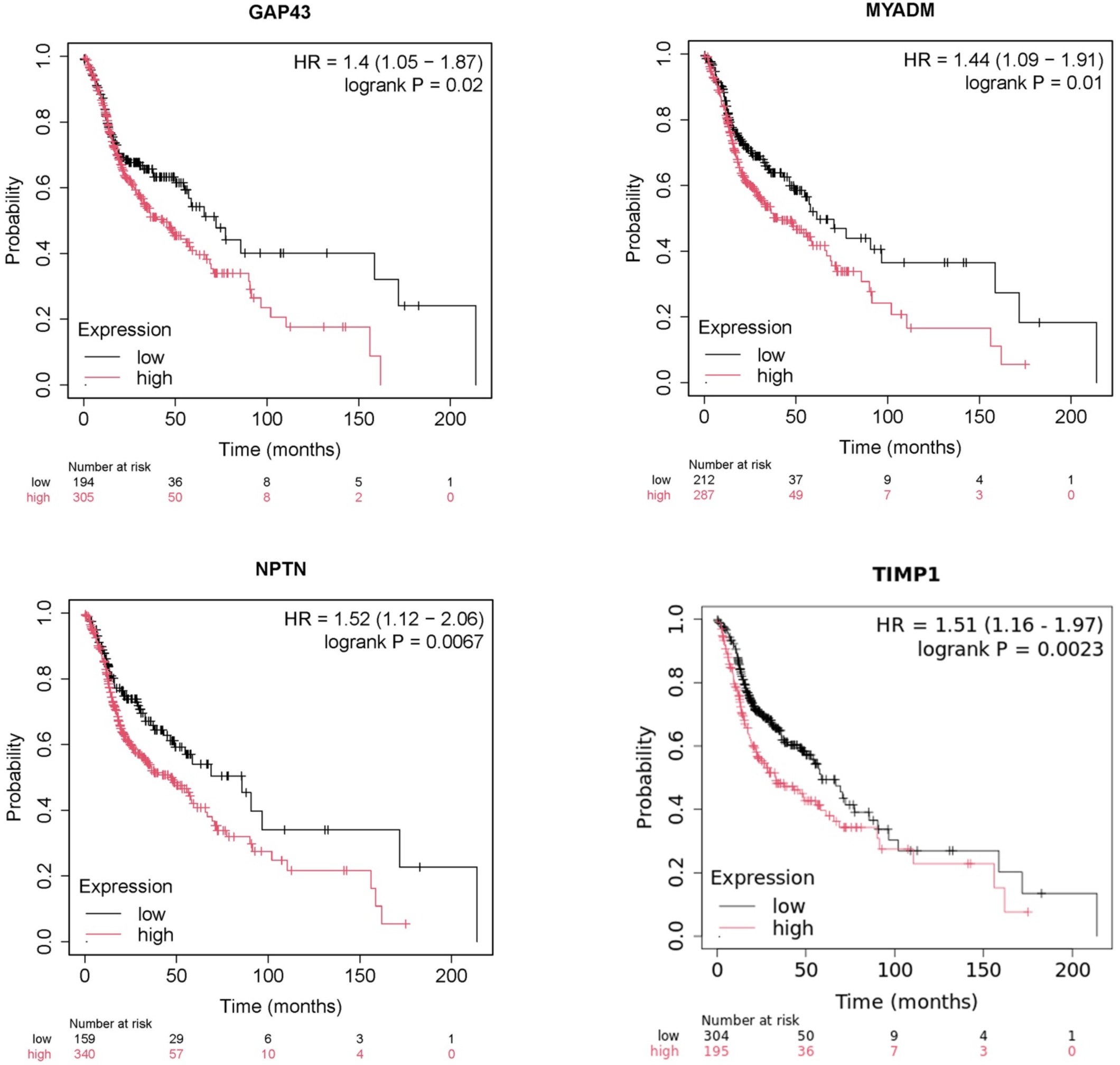
Survival plots for patients with tumors expressing high or low levels of potential EV-based biomarker candidates. Kaplan-Meier survival curves depicting the overall survival of HNSCC patients stratified into high-and low-expression groups for GAP43, MYADM, NPTN, and TIMP1. The number of patients in each group is listed under the plots. HR=Hazard Ratio. Logrank p indicates statistical significance. Data from TCGA RNA Seq for 499 patients.

## Notes

### Competing Interest Statement

The authors have declared no competing interest.

